# Genome Wide Study of Tardive Dyskinesia in Schizophrenia

**DOI:** 10.1101/386227

**Authors:** Max Lam, Keane Lim, Jenny Tay, Nina Karlsson, Smita N Deshpande, BK Thelma, Norio Ozaki, Toshiya Inada, Kang Sim, Siow-Ann Chong, Jianjun Liu, Jimmy Lee

## Abstract

Tardive dyskinesia (TD) is a severe condition characterized by repetitive involuntary movement of orofacial regions and extremities. Patients treated with antipsychotics typically present with TD symptomatology. Here, we conducted the largest GWAS of TD to date, by meta-analyzing samples of East-Asian, European, and African-American ancestry, followed by analyses of biological pathways and polygenic risk with related phenotypes. We identified a novel locus and three suggestive loci, implicating immune-related pathways. Through integrating trans-ethnic fine-mapping, we identified putative credible causal variants for three of the loci. Multivariate analyses of polygenic risk for TD supports the genetic susceptibility of TD, with relatively lower allele frequencies variants being associated with TD, beyond that of antipsychotic medication. Together, these findings provide new insights into the genetic architecture and biology of TD.

## Introduction

Tardive Dyskinesia (TD) is a persistent and potentially debilitating involuntary movement disorder characterized by choreiform, athetoid, and or dystonic movements.^1,2^ TD is largely caused by antipsychotic treatment and was first described in 1957.^3^ Although commonly observed in patients with schizophrenia, TD can occur in individuals with other psychiatric disorders, as long as they have been similarly exposed to prolonged antipsychotic treatment. The prevalence of TD in schizophrenia is estimated to be between 20-30%, and rates of TD have reduced with prescription of atypical antipsychotics.^4,5^ Nevertheless, atypical antipsychotics still carry with them a risk of developing TD, suggesting a genetic component. TD remains a clinically relevant phenotype as it has been associated with more severe schizophrenia illness, cognitive impairments, lowered quality of life and increased mortality.^6–9^

The pathophysiology of TD is currently unknown. Theories pertaining to dopamine receptor hypersensitivity, serotonergic dysfunction, GABA insufficiency and free radical damage have been put forth.^2,10,11^ It is postulated that TD is related to the schizophrenia disease process; recent reports have linked basal ganglia volume reductions to TD schizophrenia.^12^ As TD is potentially irreversible with no effective treatment, there needs to be an added emphasis on the prevention and identification of genetic risk factors. Several genes (e.g. DRD2, DRD3, MnSOD, CYP2D6, GRIN2A, GRIN2B) have been implicated in candidate gene studies, but replication remains equivoca.l^13–17^ Genome Wide Association Studies (GWAS) performed on the TD phenotype suggested that the GLI family zinc finger 2 (GLI2), heparan sulfate proteoglycan 2 (HSPG2), dipeptidyl-peptidase 6 (DPP6) and GABA pathway genes could putatively be considered susceptibility genes for TD.^18–22^ Nonetheless, these studies are limited by small samples and require further replication. Here, we report the findings of the largest GWAS of TD to date. We identified a novel locus at 16q24.1 (P = 3.01 × 10^−8^) and three suggestive loci (1p36.22, 6q23.2, 12q13.13; P < 5×10^−7^) that were associated with TD.

## Methodology

Participants from the Singapore Clinical and Translational Research program in Singapore, and the Clinical Antipsychotic Trials of Intervention Effectiveness (CATIE)^23–25^ were included in the current report. All participants met criteria for DSM-IV diagnosis for schizophrenia. Tardive dyskinesia was ascertained via the Abnormal Involuntary Movement Scale (AIMS).^26,27^ After quality control procedures, 1406 participants (280 with TD), and 6,291,020 SNPs remained. Linear Mixed Models GWA was performed via GEMMA,^28^ and independent cohorts across the two studies were meta-analyzed via fixed effects inverse variance approach in METAL.^29^ GWAS summary statistics were then subjected to functional annotation for GWAS identified loci,^30^ eQTL lookups, pathway analysis^31^ and transcriptome-wide analysis^32^ were conducted to characterize potential biological mechanisms underlying TD. Further fine-mapping analysis was also carried out to identify credible causal variants.^33–35^ A series of polygenic risk score analyses were performed to evaluate the clinical utility of measuring polygenic risk scores in TD.^36^ Finally, variants previously associated with TD were compared with the current GWAS results.^10,17,21,22,37–45^ Detailed methodological approaches are further reported in the Supplementary Information included with the current report.

## Results

### Demographics and Assessment of Tardive Dyskinesia

Demographics are reported in Supplementary Table 1. There were 71.1% males and 28.9% females with TD. There was no significant difference in gender proportion between individuals with TD and those without, χ^2^ (1, n= 1406) = .142, P = .706. There were significant differences in age, t(1404) = −14.06, P = 4.0 × 10^−42^, age of illness onset, t(392.7) = −3.06, P = 0.0024, duration of illness, t(1390) = −11.28, P = 2.76 × 10^−28^, and antipsychotic dose measured in CPZ equivalent, t(516.3) = 3.73, P = 0.002 between individuals with and without TD. These differences in demographics and clinical characteristics were further modelled using a polygenic risk score approach that allows us to examine genetic effects for TD alongside demographics effects; results are reported in subsequent sections.

**Table 1.** Top SNPs associated with tardive dyskinesia

| SNP | Symbol | Location | Position | Variant | Effect allele | Other allele | EAF | Beta | SE | P-value | HetP |
| --- | --- | --- | --- | --- | --- | --- | --- | --- | --- | --- | --- |
| rs11639774 | GSE1 | 16q24.1 | 85221868 | Intergenic | A | G | 0.168 | 0.151 | 0.0273 | 3.01E-08 | 0.067 |
| rs499646 | TNFRSF1B | 1p36.22 | 12239089 | Intronic | A | G | 0.12 | 0.1949 | 0.0364 | 8.30E-08 | 0.246 |
| rs4237808 | CALCOCO1 | 12q13.13 | 54093797 | Intergenic | T | C | 0.4914 | 0.0765 | 0.0144 | 1.08E-07 | 0.506 |
| rs6926250 | EPB41L2 | 6q23.2 | 131302473 | Intronic | T | C | 0.6872 | -0.0968 | 0.0188 | 2.54E-07 | 0.447 |

*Note.* SNP, single-nucleotide polymorphism; EAF, effect allele frequency; SE, standard error; HetP, heterogeneity P-value.

**Table 2.**
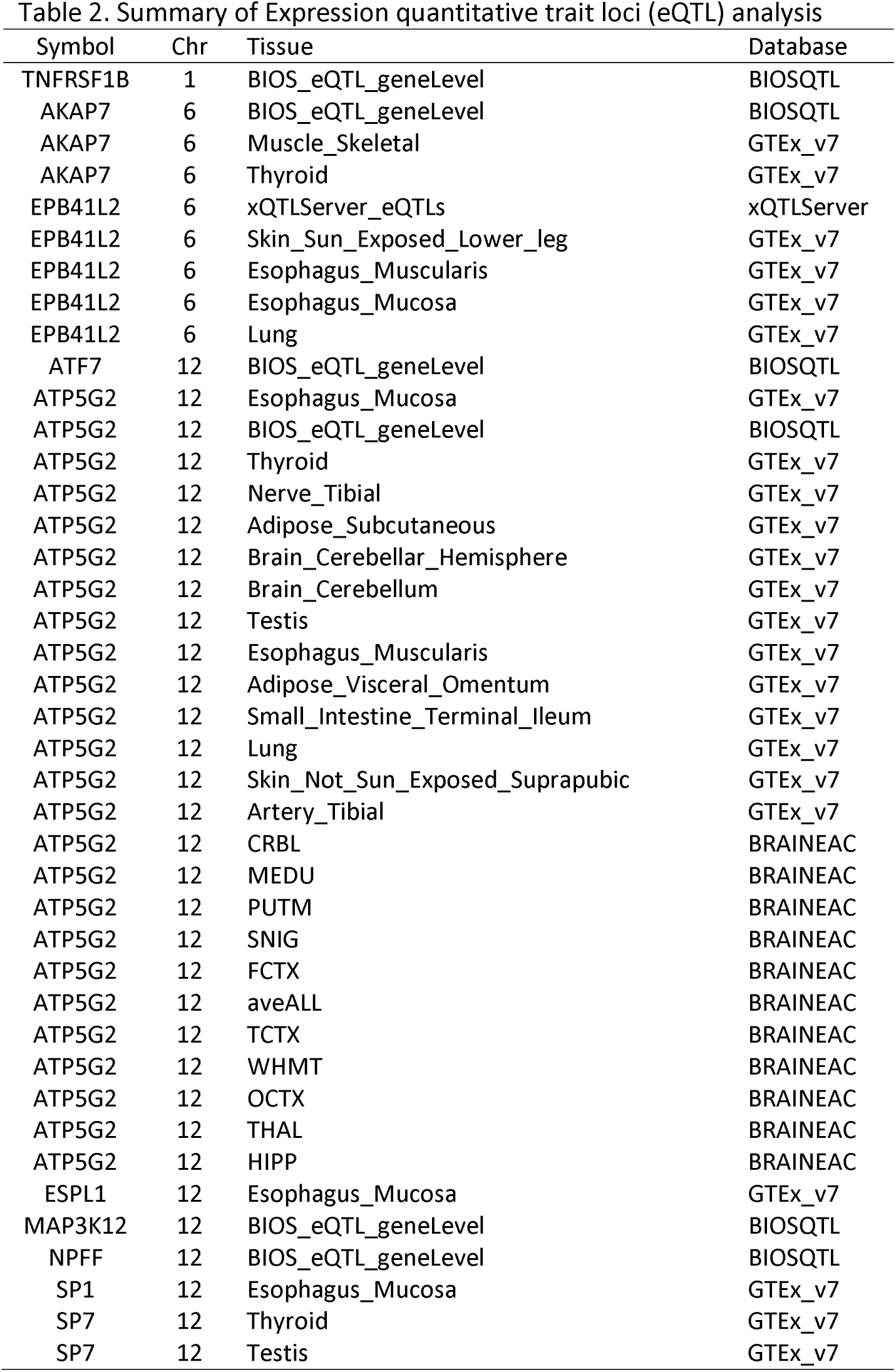
Summary of Expression quantitative trait loci (eQTL) analysis

### Genome-wide Association of Tardive Dyskinesia

Standard GWAS quality control procedures were carried out (Supplementary Figs. 1-5). Linear mixed models conducted via the GEMMA^28^ package revealed significant genome-wide association of TD at the level of the CATIE cohorts, but not the STCRP cohort (Supplementary Fig. 6). Subsequent fixed-effect inverse variance meta-analysis^29^ (λ_GC_ = 1.02; Table 1, Figs. 1–3) across the STCRP and CATIE cohorts revealed one novel locus on chromosome 16 (rs11639774, downstream of GSE1) (P = 3.01 × 10^−8^). Three other suggestive independent genomic loci (P < 5×10^−7^) were identified on chromosome 1 (rs499646, P = 8.30 × 10^−8^, TNFRSF1B), chromosome 6 (rs6926250, P = 2.54 × 10^−7^, EPB41L2) and chromosome 12 (rs4237808, P = 1.08 × 10^−7^, CALCOCO1). Due to low minor allele frequencies in the STCRP sample, two of the SNPs were only present in the CATIE cohorts. Proxy SNPs in LD within the STCRP cohort were identified using the SNP Annotation and Proxy Search (SNAP).^46^ Further meta-analysis of association results from both proxy SNPs were carried out using Fisher’s p-value meta-analysis approach (https://CRAN.R-project.org/package=metap). Fisher’s meta-analysis for the top SNP (rs11639774) and proxy SNP (rs9928615) in the STCRP cohort was significant (χ^2^(4)=40.28, P = 3.79 × 10^−08^). Similarly, results for SNP (rs499626) and proxy SNP (rs11569835) in the STCRP cohort reached suggestive significance (χ^2^(4)=36.48, P = 2.31 × 10^−07^), with both primary variants with proxy SNP meta-analysis showing consistent effects (Supplementary Table 2). GWAS markers were further annotated via ANNOVAR, eQTL, Chromatin Interaction modules within the Functional Mapping and Annotation of Genome-Wide Association Studies (FUMA)^30^ (Supplementary Tables 3, 4; Supplementary Fig 7).

**Figure 1.**
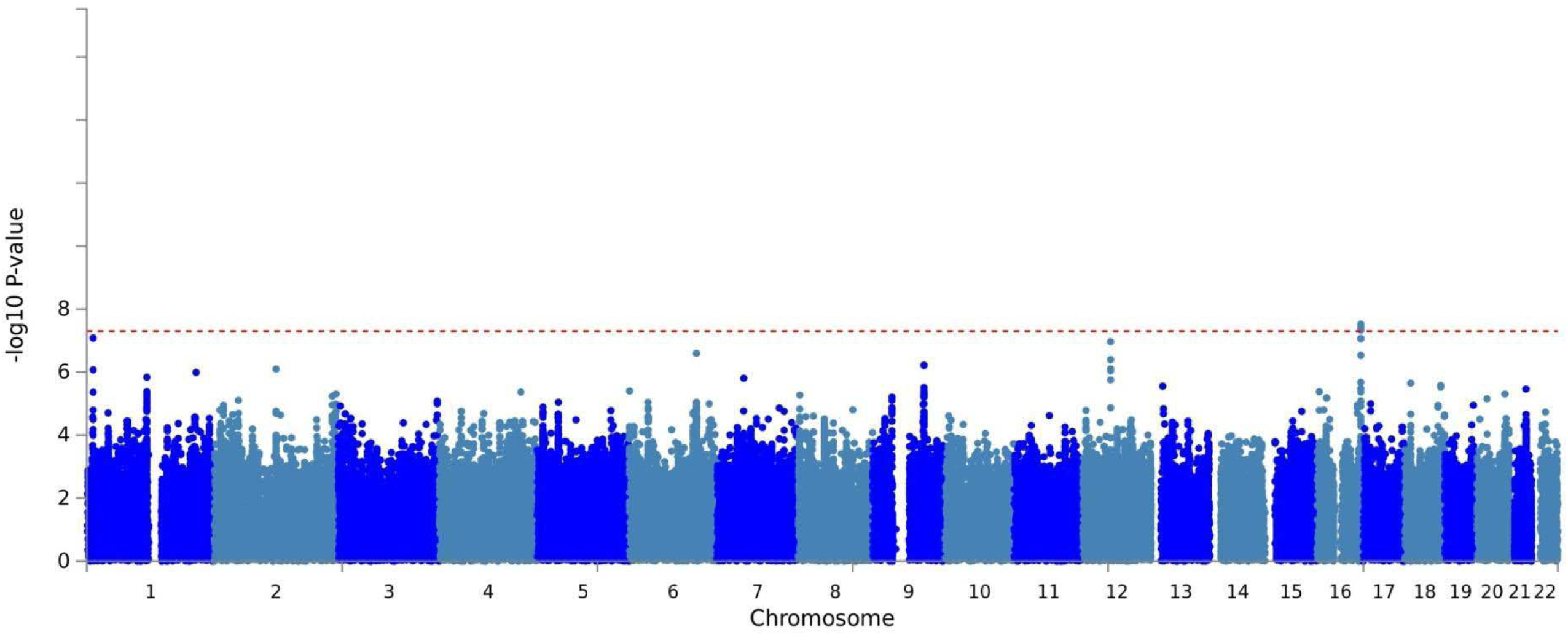
Manhattan plot for meta-analysis of STCRP, CATIE-EUR and CATIE-AFR

**Figure 2.**
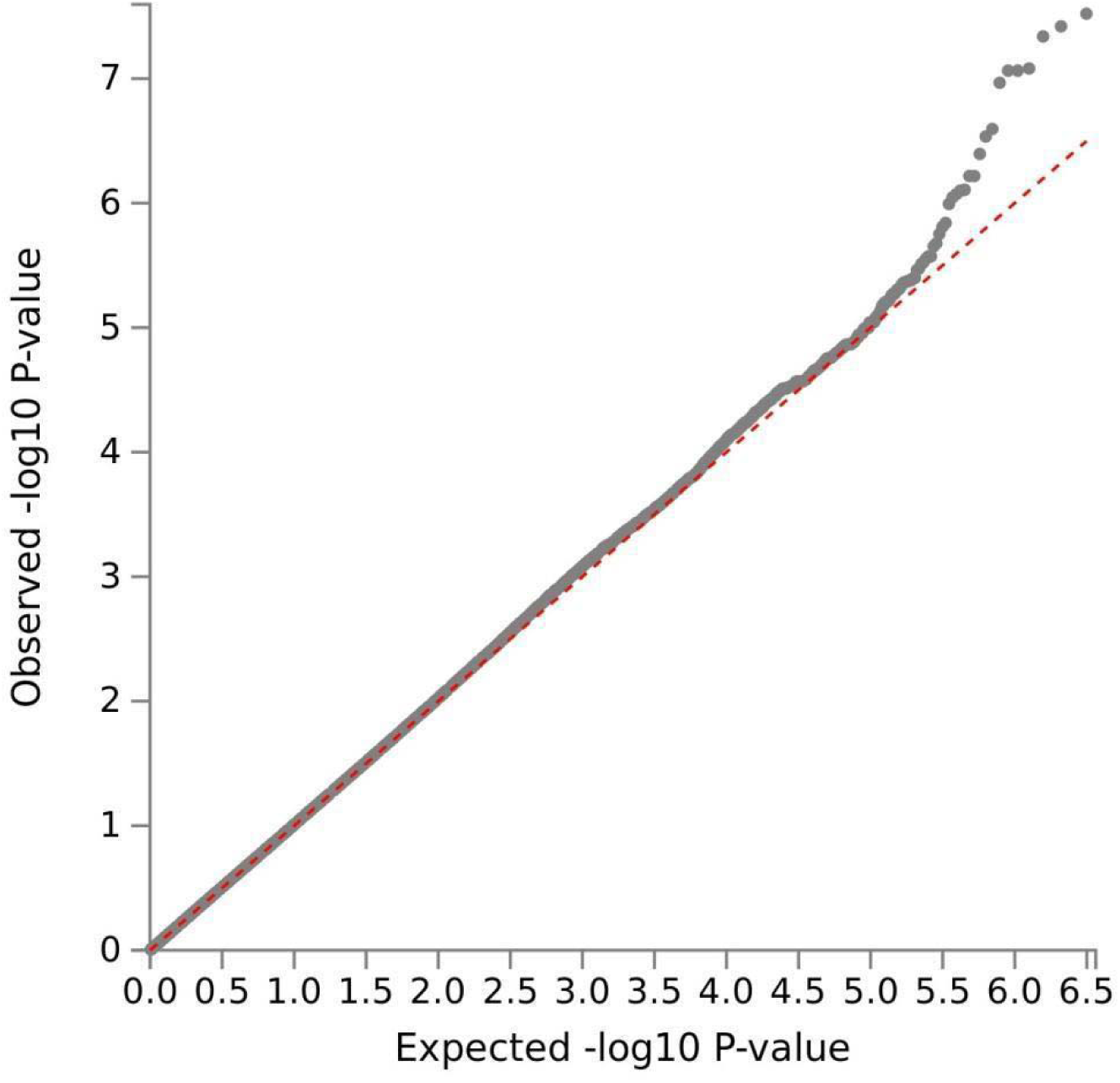
QQ plot for meta-analysis of STCRP, CATIE-EUR and CATIE-AFR. Lambda=1.02.

### Gene-based and Pathway Analysis

MAGMA was used to conduct both the gene-based and gene-set analysis.^31^ Gene-based analysis tests for association of TD with markers within the gene by mapping the SNPs to gene level, while gene-set analysis aggregates individual genes to a collection of genes with overlapping characteristics.^28^ Results of MAGMA gene-based analysis on 18,259 mapped autosomal genes after Bonferroni correction are presented in Supplementary Table 5. None of the genes were significant after Bonferroni correction. MAGMA competitive gene-set analysis^31^ was conducted using the most recent Molecular Signature Database version 6.1.^47^ Although none of the 17,199 gene-sets survived Bonferroni correction, top pathways implicated regulation of immunoglobulin production and immunoglobulin isotype switching, expression of chemokine receptors, and regulation of cell growth, and monocytes (Supplementary Table 6).

### eQTL Lookups/Transcriptome-wide Analysis

Expression quantitative trait loci (eQTL) were performed as part of FUMA^30^ GWAS pipeline. Notably, eQTL effect of ATP5G2 (Bonferroni corrected P = 1.24 × 10^−14^, Supplementary Table 7) and MAP3K12 (Bonferroni corrected p = 6.29×10^−10^, Supplementary Table 7) was observed for subthreshold genomic significance loci for 12q13.13, TNFRS1B (Bonferroni corrected p = 4.60×10^−08^, Supplementary Table 7) for 1p36.22, EPB41L2 (Bonferroni corrected p = 2.86×10^−07^, Supplementary Table 7) and AKAP7 (Bonferroni corrected p = 1.48×10^−06^, Supplementary Table 7) for 6q23.2, with expression in various tissues, including those related to motor functions (Table 2).

Transcriptome-wide analysis was implemented via MetaXcan.^32^ Unlike eQTL lookups, the MetaXcan approach further incorporates evidence from GWAS summary statistics with genome-wide gene expression profiles from the GTexV7^48^ database. This provides information of potential functional enrichment within a particular genomic locus. Here, we found lower expression of a transcript, EPB41L2, in the esophagus muscularis at Bonferroni-corrected significance (P = 0.02, Supplementary Table 8).

### Fine-mapping Analysis

Trans-ethnic fine-mapping was performed on PAINTOR v3.1^33,34^ to identify putative causal variants on the 4 loci (1p36.22, 6q23.2, 12q13.13, 16q24.1). The 99% cumulative posterior probability identified a total of 231 putative credible causal SNPs across three loci (1p36.22, 12q13.13, 16q24.1; Supplementary Table 9, Supplementary Fig. 8). From these, highly credible SNPs were defined as posterior probability > 0.8. This identified a putative causal variant for each loci 1p36.22 (rs499646, posterior probability = 0.99), 12q13.13 (rs4237808, posterior probability = 0.863), and 16q24.1 (rs28468398, posterior probability = 0.952). Notably, for loci on 1p36.22 and 12q13.13, the index SNP identified in GWAS is also a putative causal SNP identified by PAINTOR (See Figure 3). Further annotations via the Variant Effect Predictor^49^ revealed that i) rs499646 is an intronic variant 3500 bp from the promotor of TNFRSF1B and is a known protein coding variant and appears to be a loss-of-function intolerant variant ii) rs4237808 is an intergenic variant between ATP5G2 and CALCOCO1, but lies within 1000bp of two CTCF binding sites, and 3200bp from a known promotor flank iii) rs28468398 is a regulatory region variant, which lies within a transcription factor binding site on CTC-786C10.1/GSE1 gene. Notably, additional joint finemap annotations revealed that all three variants were enriched by known brain level gene expression sites close by (See Figure 3). GCTA-COJO^35^ revealed no further signals present in the loci beyond independent variants identified via LD clumping or finemapping (Supplementary Fig. 9).

**Figure 3.**
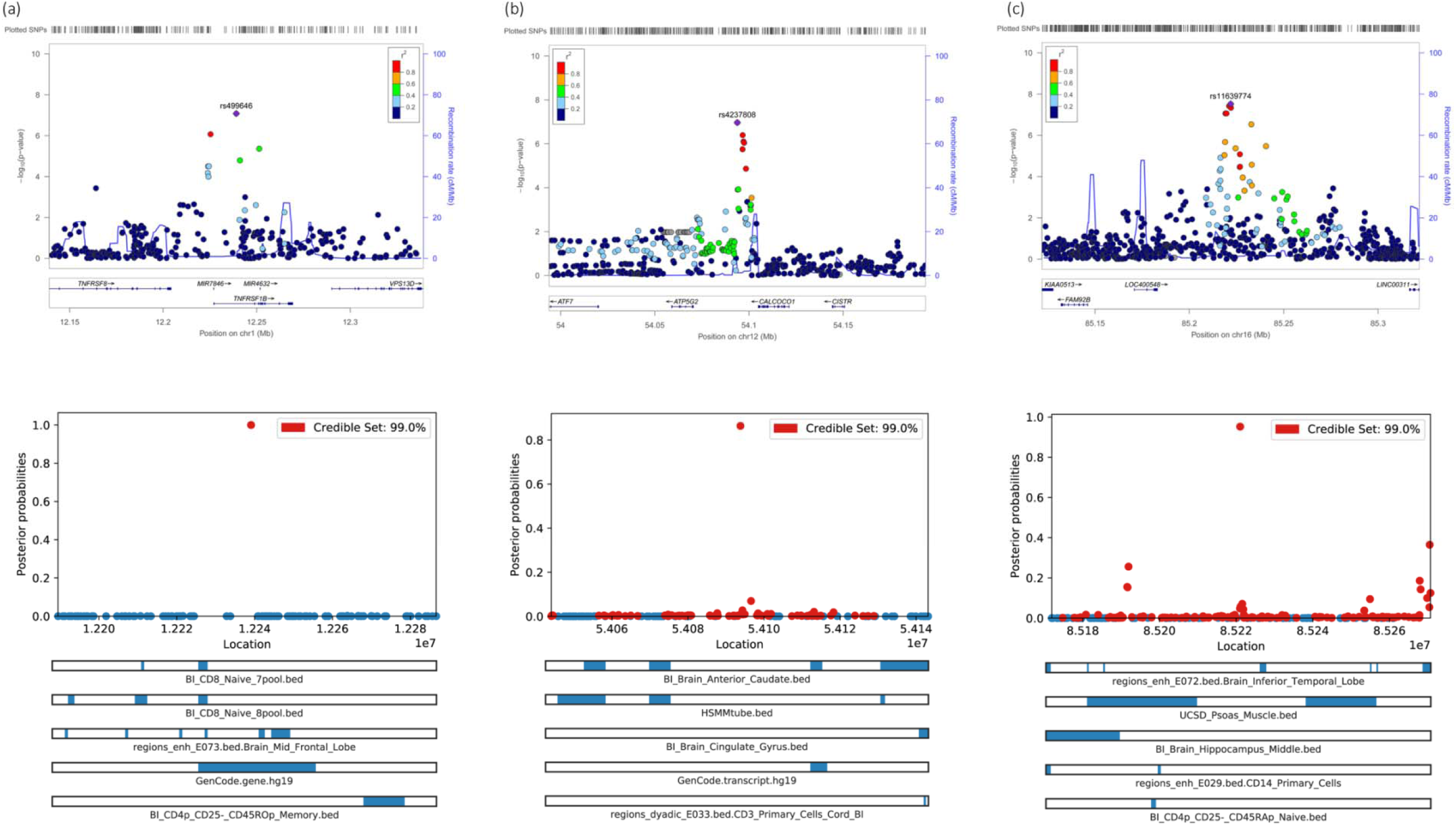
Regional loci and fine-mapping plots Top panel represents regional plots for (a) Chromosome 1, (b) Chromosome 12, and (c) Chromosome 16. Bottom panel represents the visualization of 99% credible SNP set for (a) Chromosome 1, (b) Chromosome 12, and (c) Chromosome 16, against location of SNP, with top annotation bars.

### Polygenic Risk Modelling of TD with Other Diseases and Traits

Polygenic risk scores modelling via PRSice2^36^ (Supplementary Figs. 10) revealed best polygenic association of TD with amyotrophic lateral Sclerosis (ALS), albeit only two threshold remained significant after Bonferroni correction (P_T_ = 0.5; P_T_ = 1.0). All other traits tested did not yield significant polygenic risk models after Bonferroni correction. Further analysis of ALS PRS-based pathway analysis indicated “misfolded protein” as a top pathway shared between TD and ALS (Supplementary Fig. 11), though, this remains a trend finding.

### Risk Prediction of TD with PRS and Clinical Variables

Additionally, we investigated if TD was associated as a function of rare or common alleles. Allele frequency from the 1000 genomes phase 3 reference panel were entered as weights for polygenic risk score (PRS) calculation. Lower PRS scores indicated presence of low frequency allele variants while higher scores represent higher concentration of common variants in the genome. Partial correlation between TD and clinical variables are presented in Supplementary Table 10. Multivariate logistic regression modelling revealed that polygenic risk scores (β = −1.437, SE = 0.277, P = 2.22×10^−07^), age (β = 0.086, SE = 0.008, P = 5.64×10^−28^) and severity of psychopathology (PANSS total score) (β = 0.012, SE = 0.004, P = 0.004) significantly predicted TD ( χ^2^ (10, N = 1214) = 222.54, P = 3.14×10^−42^, Nagelkerke R^2^ = 0.263). The strongest predictor of TD was the allele frequency weighted polygenic risk score, with an odds ratio of 4.21 (95% CI = 2.44-7.25) after accounting for covariates in the model. The alternative hypothesis was investigated by not weighting PRS by population allele frequency, computing summation of genome-wide genotype counts per individual. No significant association was found (Supplementary Table 10), indicating TD may be associated with variants of lower allele frequencies.

### Risk Prediction of TD with PRS and Clinical Variables – Subtyped by Medicatio

To further investigate the effect of medication type on TD status, three models of multivariate logistic regression (i.e., Individuals taking typical antipsychotics only, atypical antipsychotics only, and combination of typical and atypical antipsychotics) were conducted on variables that showed association with TD (Supplementary Table 10). In the typical antipsychotics only model (Nagelkerke R^2^ = 0.208), significant association was found for age (β = 0.085, SE = 0.011, P =1.56×10^−15^) and PANSS total (β = 0.014, SE = 0.007, P = 0.036), and marginal association was found for allele frequency weighted PRS (β = −0.673, SE = 0.347, P = 0.052). However, only age remained significant after Bonferroni correction (P = 0.05/3). In the atypical antipsychotics model (Nagelkerke R^2^ = 0.186), only age (β = 0.081, SE = 0.016, P = 5.17×10^−07^) and allele frequency weight PRS (β = −2.751, SE = 0.555, P = 7.12×10^−07^) were associated with TD status. Both age and allele frequency weighted PRS remained significant after Bonferroni correction. In the atypical and typical antipsychotic combined model (Nagelkerke R^2^ = 0.271), age (β = 0.134, SE = 0.037, P = 2.86×10^−4^), allele frequency weighted PRS (β = −4.224, SE = 1.47, P = 0.004), and gender (β = 1.686, SE = 0.852, P = 0.048). Only age and PRS remained significant after Bonferroni correction. Results suggest that polygenic score might be related to prescription of medication type. To investigate this more directly, the sample is further stratified via polygenic scores, comparing the top (high polygenic risk group) and bottom 10 (low polygenic risk group) percentiles of the sample for the association between TD (Controls v Case) x Medication Type (Typical v Atypical Antipsychotics). No significant differences were observed in the high polygenic risk score group (Typical*Controls = 37.9%; Atypical*Case = 24.1%; Typical*Case = 19.8%; Atypical*Case = 12.1%; OR = 1.05; SE = 2.25; P = .915). In the low polygenic risk score group, we found significant differences in the proportions of individuals, taking typical antipsychotics and having TD compared to those taking atypical antipsychotics (Typical*Controls = 38.5%; Atypical*Controls = 36.9%; Typical*Cases = 9.8%; Atypical*Cases = 0.8%; OR = 11.49; SE = 5.71; P = 0.021).

### Look-up of Past Tardive Dyskinesia Studies

Variants previously found to be associated with TD were meta-analyzed with results from the current GWAS (Supplementary Table 11).^10,17,21,22,37–45^ The following variants were replicated in the current study CYP1A2 (N = 1725, P = 0.018), DRD1 (N = 1788, P = 0.045), GRIN2A (N = 1837, P = 0.012), GRIN2B (N = 1057, P = 0.0018), HSPG2 (N = 1572, P = 0.0051), and DPP6 (N = 1701, P = 0.00068). These genes implicated pathways involving drug metabolism, dopamine, and glutamate.

## Discussion

To our knowledge, here we report the largest GWAS for TD. A single novel locus at 16q24.1 was found to predict TD. We also identified 3 suggestive loci (1p36.22, 6q23.2, 12q13.13) that were associated with TD. Of these, putative causal variants were identified for three of the loci. The top GWAS hit at 16q24.1, in the GSE1 coiled coil protein gene, encodes for a proline rich protein which was reported to be a subunit of BRAF35-HDAC (BHC) histone deacetylase complex.^50^ This gene has been known to be implicated in the proliferation, migration, and invasion of breast cancer cells.^51^ A look-up in GWAS catalog revealed GSE1 was also associated at GWAS significance with platelet count and distribution,^52^ and suggestive GWAS significance with amyotrophic lateral sclerosis^53^ and sulfasalazine-induced agranulocytosis.^54^ GSE1 contributes to a gene-set that are predicted targets of a microRNA biomarker for schizophrenia.^55^

Other suggestive associations with TD included TNFRSF1B (1p36.22), EPB41L2 (6q23.2), and CALCOCO1 (12q13.13). The tumor necrosis factor receptor superfamily member 1B (TNFRSF1B) is a protein coding gene that mediates anti-apoptotic signaling. Expression of TNFRSF1B is specific to cells in the immune systems, specific neuronal subtypes, certain T-cells subtypes, and endothelial cells.^56^ This appears to be supported by results from the competitive gene-set enrichment analysis, albeit non-significant, revealed top pathways that implicated immunoglobulin production, chemokine receptors and monocytes. These findings appear to support existing theories on the role of immune and inflammation in the pathogenesis of TD.^10,11^.

The erythrocyte membrane protein band 4.1 like 2 (EPB41L2) is involved in actin and cytoskeletal protein binding. More recently, EPB41L2 deficiency was found to result in myelination abnormalities in the peripheral nervous system, leading to motor neuropathy in a mice study.^57^ This putative association of EPB41L2 deficiency is consistent with the direction of effect found in our GWAS results, suggesting that the expression of EPB41L2 might confer a protective effect against motor neuropathy. Functional enrichment in and around the EPB41L2 based on the MetaXcan finding is intriguing, further research is needed to dissect the function of EPB41L2 in TD pathophysiology.

The calcium binding and coiled-coil domain 1 (CALCOCO1) is a protein coding gene that is involved in the activation of transcriptional activities of targets genes in the Wnt signaling pathway, neuronal receptor and aryl hydrocarbon receptor (AhR).^58^ Notably, one function of the AhR, a ligand-based transcription factor, is the regulation of transcriptional activity for drug metabolizing enzymes, including the family of cytochrome P450 (CYP) genes.^59^ The family of CYP has been postulated to be a candidate for TD susceptibility.^10^ Specifically, CYP enzymes such as CYP1A2 metabolizes antipsychotics (e.g., clozapine, olanzapine, and haloperidol) and antidepressants (e.g., fluvoxamine).^59^ Meta-analysis of CYP1A2 from past and current study revealed that this gene was significantly associated with TD at trend level (P < 0.05; Supplementary Table 11).

Polygenic risk score analysis on traits with polygenic architecture revealed strongest overlapping genetic polygenic risk of TD and ALS. However, the lack of robust genetic overlap between TD and other seemingly related traits, such as neurodegenerative conditions and autoimmune conditions does not preclude the plausible shared genetic architecture of these traits. Rather, ancestry difference in LD patterns and allele frequencies of the target and training dataset could result in the poorer polygenic prediction; the summary statistics of these traits were primarily of European ancestry. Other factors including the lack of power, use of different reference panel (i.e. HapMap and 1000 genomes project) in the various traits could plausibly contribute to the lack of robust PRS findings. Nevertheless, the shared protein misfolding pathway between TD and ALS appears to reconcile the findings from our MAGMA pathway analysis. Proteopathies commonly implicated in neurodegenerative conditions have been proposed to also underlie TD.^60,61^ Reports implicated the accumulative role of proteopathies in neurotoxicity, synaptic dysfunction, and its bidirectional effect with oxidative stress, neuroinflammation, and its consequence on the immune system.^60,61^ However, PRS-based pathway analysis currently implemented in PRSice2 is performed under a self-contained model; hence, further replication of these results is needed.

Multivariate analyses of PRS and clinical variables, grouped by medication type, appear to support the increased risk of TD with typical antipsychotics. Traditionally, antipsychotics, particularly typical antipsychotics, have been thought be a cause of TD due to the blockade of dopamine receptor,^62^ and the expression of D2 receptors mainly in the basal ganglia.^63^ Here, we present evidence that show in part that individuals with low polygenic risk to TD are in fact more likely to eventually suffer from TD after being prescribed typical antipsychotic medication compared to individuals prescribed atypical antipsychotic. On the other hand, if one has a high polygenic risk to TD, we demonstrate that there is potentially an equal probability of having TD regardless of the type of antipsychotic prescribed. Though the sample sizes that drive these findings are relatively small and in need of further replication, we provide the first evidence in psychiatric GWAS the utility of polygenic risk score being a screener in clinical practice. We also acknowledge that longitudinal information in subsequent studies might be necessary to identify cumulative drug effects and their interaction with the genetic architecture underlying TD. Nevertheless, taken together, these results appear offer prima facie evidence to the earlier understanding of TD etiology; but also point to more complex biological interactions between the pharmacodynamics of antipsychotics, and the genetic predisposition to TD. Further work is necessary to understand the phenomenon elucidated in the current report.

Emergent results from the current report demonstrate the polygenic architecture of TD. While the current study remains underpowered for GWAS analysis (Supplementary Fig. 12), we present the first and largest GWAS of TD to date. The findings reported here suggest that multiple overlapping biological systems might contribute to the etiopathogenesis of the condition and in some, exacerbated by pharmacological compounds administered for the treatment schizophrenia symptomatology. Taken together, results suggest that TD is associated with significant proteopathies, disrupted neuronal function within the brain and potentially including muscular innervation, existing in a background of inflammation. Further work is necessary to identify the cascade of biological disruptions and events that trigger TD symptomatologies. Further work is needed to further unravel the biology of these risk variants and pathways identified, which could potentially inform the development of therapeutic targets for TD and guide treatment decisions in the clinic.

## Acknowledgements

This research is supported by grants from the Ministry of Health Singapore, National Medical Research Council (Grant No.: NMRC/TCR/003/2008, NMRC/CG/004/2013). ML is supported by the National Medical Research Council Research Training Fellowship (Grant No.: MH095:003/008-1014). The authors thank the NIH for providing limited access datasets for the NIMH CATIE (ClinicalTrials.gov identifier NCT00014001, NIMH contract #N01MH90001). We would like to express our gratitude to Professor Arinami Tadao for his valuable comments. The authors also thank all participants and researchers who contributed to the collection of this data.

## Author Contributions

ML and KL ran the analysis and drafted the manuscript, JJL and JL designed the study and supervised preparation of the manuscript. All other authors were involved in the drafting and approved the final manuscript. JL takes responsibility for data access.

## Web Resources

FUMA, http://fuma.ctglab.nl/

GAS Power Calculator, http://csg.sph.umich.edu/abecasis/cats/gas_power_calculator/index.html

GCTA, https://cnsgenomics.com/software/gcta/

GEMMA, http://www.xzlab.org/software.html

GWASCatalog, https://www.ebi.ac.uk/gwas

LD-HUB, http://ldsc.broadinstitute.org/

MAGMA, https://ctg.cncr.nl/software/magma

METAL, https://genome.sph.umich.edu/wiki/METAL_Documentation

MetaXcan, https://github.com/hakyimlab/MetaXcan

Michigan Imputation Server, https://imputationserver.sph.umich.edu/index.html

MSigDB, http://software.broadinstitute.org/gsea/msigdb

PAINTOR, https://github.com/gkichaev/PAINTOR_V3.0

PLINK1.9, https://www.cog-genomics.org/plink2

PRSice2, https://github.com/choishingwan/PRSice/wiki

PredictDB, http://predictdb.org/

R, https://www.r-project.org/

SNAP, http://www.broad.mit.edu/mpg/snap/

Variant Effect Predictor, https://www.ensembl.org/vep

